# The phoenix hypothesis of speciation

**DOI:** 10.1101/2022.06.16.496444

**Authors:** Ryo Yamaguchi, Bryn Wiley, Sarah P. Otto

**Author notes:** Corresponding author: Ryo Yamaguchi.

## Abstract

Genetic divergence among allopatric populations builds reproductive isolation over time and is thought to be the major mechanism underlying the formation of new species. This process is accelerated when populations face a changing environment, but abrupt change also places populations at risk of extinction. Here we use simulations of Fisher’s geometric model with explicit population dynamics to explore the genetic changes that occur in the face of extreme environmental changes to which populations must adapt or go extinct. We show that evolutionary rescue leads to the fixation of mutations whose effects are larger on average and that these mutations are more likely to lead to reproductive isolation, compared with populations not at risk of extinction. We refer to the formation of new species from the ashes of populations in decline as the phoenix hypothesis of speciation. The phoenix hypothesis predicts more substantial hybrid fitness breakdown among populations surviving a higher extinction risk. The hypothesis was supported when many loci underlie adaptation. When, however, there was only a small number of potential rescue mutations, we found that mutations fixed in different populations were more likely to be identical, with parallel changes reducing isolation. With a limited genomic potential for adaptation, we find support for a modified version of the phoenix hypothesis where reproductive isolation builds fastest in populations subject to an intermediate extinction risk. While processes driving extinction lead to the loss of lineages with deep evolutionary histories, they may also generate new taxa, albeit taxa with minimal genetic differences.

## Introduction

Biological diversity emerges from the interplay between speciation and extinction. Historically, these two factors have been studied as independent processes, but recent speciation studies have emphasized the need to consider population persistence for new taxa (Dynesius and Jansson 2014; Harvey et al. 2019). In addition, extreme environmental changes may affect the development of reproductive isolation as well as the risk of extinction, a possibility that we explore here through simulations. Here we explore how extinction risk itself could impact the likelihood of speciation.

Recent empirical studies have suggested that adaptation to a new environment facilitates the origin of species (Schluter 2000). Yet, an abrupt environmental change poses a risk of extinction and species loss. Adaptation to a rapidly changing environment enables populations to accumulate phenotypically large-effect mutations, leading to severe fitness reduction in hybrids referred to as hybrid breakdown (Chevin et al. 2014; Yamaguchi & Otto 2020). Although large-effect mutations were seen as unlikely to contribute to adaptation because small effect changes are much more likely to be beneficial (Orr 2005a), there is a growing list of studies that have identified large effect loci contributing to adaptive divergence, particularly in response to environmental shifts, such as *Mimulus* monkeyflower (Bradshaw et al. 1998), *Peromyscus* oldfield mice (Steiner et al. 2007), maize (Doebley 2004), Ectodysplasin (Eda) allele in three spine sticklebacks (Colosimo et al. 2005; Jones et al. 2012; Schluter et al. 2021), *Heliconius* Müllerian mimicry patterns (Baxter et al. 2009), and beak shape (Alx1) and beak size (Hmga2) in Darwin’s finches (Lamichhaney et al. 2015, 2016). For populations at risk of extinction, if they fail to adapt, those populations that do survive may be especially likely to accumulate large-effect mutations (Osmond et al. 2020). Here we explore the effect of evolutionary rescue on the distributions of fixed mutations and the consequences for speciation.

Experimental evolution studies have also confirmed that large-effect mutations contribute to adaptation in a variety of species adapting to a new environment. In these studies, large-effect mutations typically fix early during adaptation, followed by a series of many smaller-effect substitutions (e.g., Holder and Bull 2001; Collins and de Meaux 2009; Estes et al. 2011). Using Chlamydomonas, Collins and de Meaux (2009) revealed that populations adapting to an abrupt environmental change showed evidence of substitutions of large effect contributing to the earlier phase of adaptation, while populations adapting to a slowly changing environment showed patterns consistent with adaptation via mutations of small effect. Following an environmental change, *Saccharomyces cerevisiae* accumulate large-effect mutations and develop strong intrinsic incompatibilities, even when adapting to similar ecological changes (Kvitek and Sherlock 2011; Ono et al. 2017). These empirical studies suggest that environmental changes are likely engines of speciation for allopatric populations.

For populations that must adapt or face extinction (“evolutionary rescue” scenario, see Bell and Gonzales (2009)), environmental stress initially causes the population to decline, which limits the potential for small-effect mutations to spread (Otto and Whitlock 1997), leading to enrichment of large-effect substitutions among populations that can adapt and persist in the new environment (Osmond et al. 2020). This enrichment of large-effect mutations may thus drive higher rates of speciation. The influence of population dynamics and extinction risk on the evolutionary rate of reproductive isolation has not, however, been explored.

We investigate the hypothesis that extinction risk may elevate the chance of speciation, which we refer to as the phoenix hypothesis of speciation. The hypothesis is named after the Greek myth of the phoenix, which burns to ashes from which the next generation emerges. The phoenix hypothesis predicts that there is a greater degree of reproductive isolation among populations experiencing extinction risk, conditional on the populations successfully adapting and persisting.

Mutations producing large phenotypic changes may, however, be more likely to fix in parallel among different populations, because there is a limited set of possible genetic pathways to adaptation (as seen empirically, e.g., Fang et al. 2021, and predicted theoretically, e.g. Orr 2005b; see also Yeaman et al. 2018). If the number of potentially beneficial mutations in a given new environment is small, the probability of evolving similar phenotypes based on shared genetic mechanisms (i.e., parallel evolution) is expected to be higher when the environment changes abruptly. Parallel adaptation interferes with the development of reproductive isolation (Thompson et al. 2019). Thus, adaptation to extreme environmental shifts is expected to involve large-effect mutations, which contribute to reproductive isolation, but also parallel mutations, which do not. It thus remains unclear what the level of incompatibility is between populations adapting to the same environmental change that do survive extinction.

Thus, depending on the extinction risk of the parent population, we have contrasting expectations for reproductive isolation, with large but more parallel mutations accumulating during evolutionary rescue. To test the phoenix hypothesis, we extend Fisher’s geometric model to simulate mutation-order speciation among diploid populations facing an extinction risk. When many genes could potentially contribute to rescue, we confirm the phoenix hypothesis and show that more reproductive isolation evolves in populations facing higher extinction risks (Figure 1a). When the number of loci available for adaptation is finite and parallel mutations are more likely, our results support a modified version of the phoenix hypothesis: a moderate extinction risk results in more extensive reproductive isolation among surviving populations (Figure 1b). When facing a moderate extinction risk, populations are more like to survive and adapt via different, large-effect mutations, generating substantial incompatibilities among populations.

**Figure 1.**
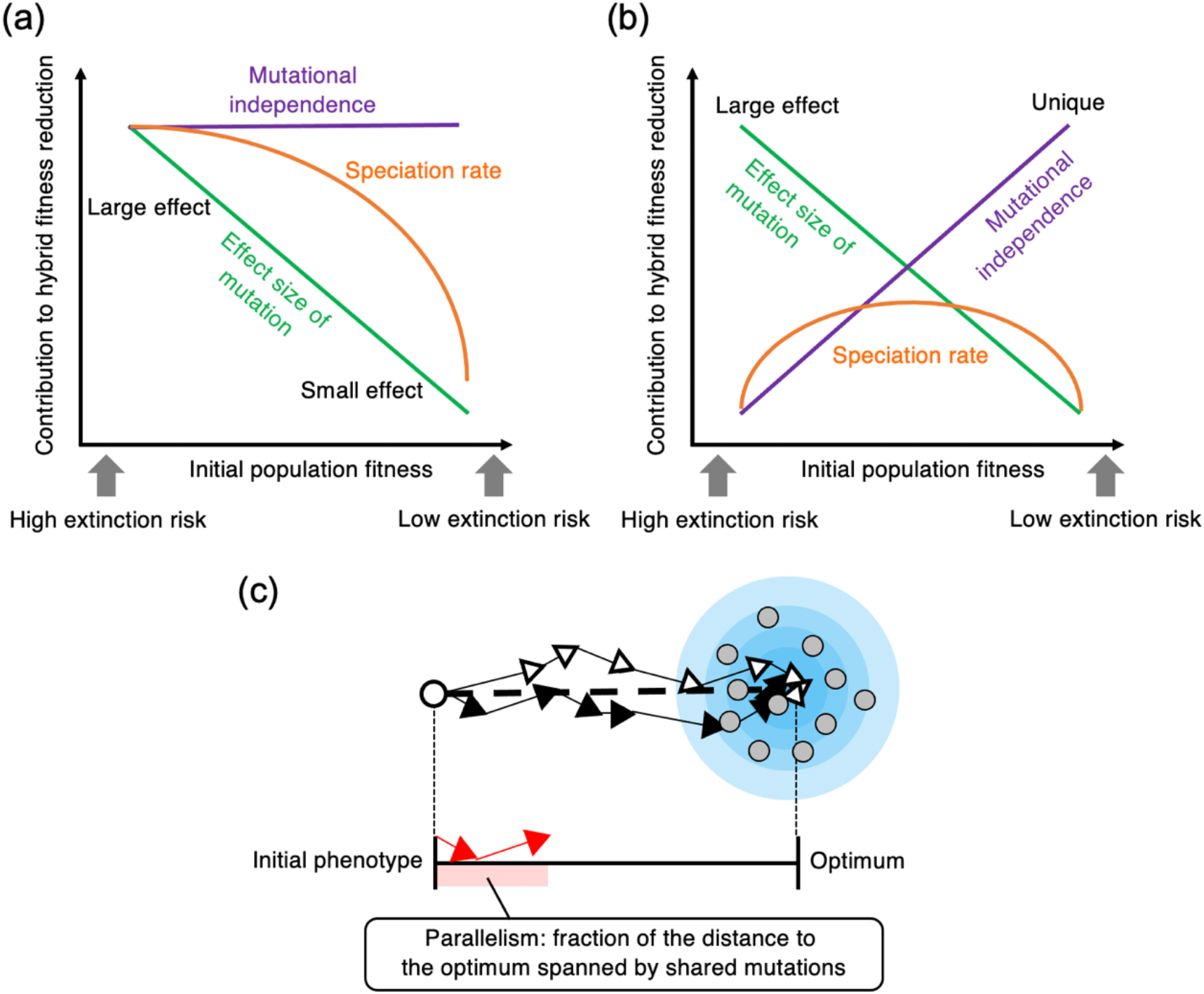
(a) The phoenix hypothesis of speciation. When extinction risk is high, large-effect mutations are likely to accumulate (green). Conversely, when extinction risk is low, small-effect mutations are likely to accumulate. Under the infinite-sites assumption, no common mutation accumulates among populations (purple). A greater degree of reproductive isolation develops under a high extinction risk (orange). (b) When the number of loci available for adaptation is finite, large-effect mutations are more likely to be common among populations (purple). Given the contrasting relationships with mutational effect sizes (green), an intermediate extinction risk can promote speciation (orange, a modified version of the phoenix hypothesis). (c) Conceptual illustration of parallelism. Here we show the case of a two-dimensional phenotype with a new optimum at the center of the blue region, illustrating the adaptive paths taken by two independent populations (black and white). A white dot and grey dots represent the position of initial phenotype and hypothetical hybrids’ phenotype, respectively. Red arrows are the mutations that are shared between the populations.

### Model

We conduct individual-based simulations of Fisher’s geometric model (FGM) with explicit population dynamics of a sexually reproducing diploid organisms with non-overlapping generations, using SLiM3.3 (Haller and Messer 2019). Table 1 summarizes the notation and gives the typical parameters used in this study.

**Table 1.** Notation and parameters used. Default values are given in square brackets, except where noted in the text. We run simulations for different values of *L* and *u* when their product (genome-wide mutation rate) is fixed as *L* × *u* = 5 × 10^−3^. Specifically, (*L, u*) = (50, 10^-4^), (100, 5 ×10^-5^) and (10000, 5 ×10^-7^).

| Notation | Description |
| --- | --- |
| $n$ | Number of phenotypes [4] |
| $\mathbf{z}_i = \{z_{i,1}, z_{i,2}, \dots, z_{i,n}\}$ | Phenotype of $i$th individual |
| $\mathbf{o} = \{o_1, o_2, \dots, o_n\}$ | Optimal phenotype ($o_1 = 2$ and otherwise 0) |
| $q$ | Parameter translating the phenotypic distance from the optimum into the strength of selection [1] |
| $k$ | Degree of epistasis [2] |
| $v$ | Mutation size, drawn from an exponential distribution with mean $\lambda = [0.4]$ |
| $\xi_j, \omega$ | Random number drawn from a standard normal distribution |
| $\Delta \mathbf{z}_i = \{\Delta z_{i,1}, \Delta z_{i,2}, \dots, \Delta z_{i,n}\}$ | Phenotypic change of $i$th individual due to mutation |
| $s$ | Average strength of selection |
| $N$ | Number of individuals |
| $N_0$ | Initial number of individuals in a population [2000] |
| $u$ | Mutation rate [$10^{-4}$, $5 \times 10^{-5}$, $5 \times 10^{-7}$] |
| $L$ | Genome size (number of sites) [50, 100, 10000] |
| $m$ | Recombination rate between consecutive sites [0.01] |
| $r$ | Intrinsic growth rate (no selection) [0.8, 1.0, 1.4] |
| $\beta$ | Constant birth and death rates [0.1] |
| $K$ | Carrying capacity [2000] |
| $c$ | Degree of pleiotropic side effect in mutation library [0, 0.5, 1.0] |

#### Fitness landscape

FGM assumes multivariate stabilizing selection around a single optimum. Similar to Yamaguchi and Otto (2020), we model this by defining the phenotype of an individual as an *n*-dimensional vector, where *n* is the number of traits relevant to adaptation in a given environment. The phenotype of the *i*th individual is denoted as ***z**_i_* = {*z*_*i*,1_,*z*_*i*,2_,…, *z_i,n_*}, where *z_i,j_* is the phenotypic value for trait *j*.

The fitness of an individual depends on the distance of its phenotypic values from the optimum ***o*** = {*o*_1_, *o*_2_,…, *o_n_*}, measured as the Euclidean distance 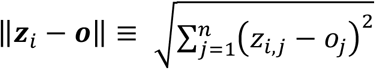. We assume that the optimal phenotype is common to all individuals and populations. An individual’s relative fitness is given by:

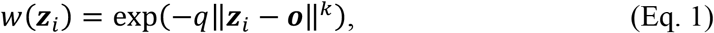

where *q* and *k* determine the decay rate and the curvature (the degree of epistasis) of the fitness landscape, respectively (Tenaillon 2014). It should be noted that isotropic selection is assumed throughout the paper; meaning that there is equally strong selection on all *n* traits at a given distance to the optimum.

#### Mutation and genetic architecture

To construct trait values from the accumulated mutations, we assume that each mutation additively contributes to the phenotypic value. Thus, an individual’s phenotype is determined by the sum of effect sizes of all mutations affecting the traits plus the original phenotype (taken to be at the origin). While each mutation has an additive effect on the phenotype, the effect converted to fitness is not additive but depends on the location of the set of phenotypic values on the fitness landscape given by Eq. 1.

For homozygous diploids, we define the effect of a mutation occurring in the *i*th individual by Δ***z_i_*** = {Δ*z*_*i*,1_, Δ*z*_*i*,2_,…, Δ*z_i,n_*}, with Δ*z_i,j_* describing the mutational effect on trait *j*. We assume that a single mutation affects each phenotype independently, with the change Δ*z_i,j_*, proportional to the value drawn from a standard normal distribution, denoted as *ξ_j_* (Hartl and Taubes 1998; Fraïsse *et al*. 2016). The overall effect of a single mutation (its length across all phenotypic axes) was set to *v*, drawn from an exponential distribution with mean *λ*.

We also consider how the degree of pleiotropic side effects impacts the likelihood of parallel adaptation and the development of reproductive isolation. To vary the strength of pleiotropy, we allowed mutations to have a main effect on only one axis (weighted by an amount *c*), as well as a randomly drawn mutation vector (weighted by 1–*c*). Specifically, for each mutation, we randomly choose a focal trait *k* and set Δ*z_i,k_* = ±*cv*. In addition, we let the mutation randomly affect each of the traits, *j* (including *k*), by an amount 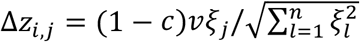. This mutation scheme is exactly the same as the original FGM when *c* = 0 (isotropic mutation), while there is no pleiotropy if *c* = 1. The phenotypic value of the mutant individual was then set to *z_i_* + Δ*z_i_*. The above describes the properties of homozygous mutations. Heterozygous diploids are assumed to be shifted half as far in phenotype space, Δ***z_i_***/2 (i.e., we assume additivity on a phenotypic scale, although dominance depends on the mutation vector and position relative to the optimum).

Mutations can occur with equal probability *u* at each locus. The toral number of loci is *L*. Recombination is modeled by assuming that cross-over events occur uniformly at random at rate *m* across loci. We prepare a mutation library to assign Δ*z_i,j_* to each locus for each potential mutation that may occur in any of the populations during a given replicate simulation. By not assuming an infinite sites model in which all novel mutations are unique, adaptive mutations can sometimes be shared between two populations evolving independently.

#### Population dynamics

We consider a population of diploid individuals with a time-dependent population size *N*(*t*). We first calculate the extinction risk analytically, modelling the population dynamics as a birth-death process with birth and death rates *b*(*t,N*(*t*)) and *d*(*t,N*(*t*)), respectively:

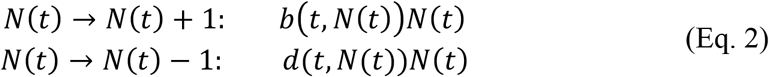

The expected change of *N*(*t*) over a small-time interval Δ*t* is E[Δ*N*|*N*(*t*)] = *r*(*t,N*(*t*))*N*(*t*)Δ*t*, where *r*(*t,N*(*t*)) = *b*(*t,N*(*t*)) = *d*(*t,N*(*t*)). Following Uecker and Hermisson (2011), we consider the process of mutation accumulation with a population that follows logistic growth (or decline) until it has reached its carrying capacity *K*. We assume that a decreasing availability of resources per individual leads to a lower birth rate and that the death rate depends on the strength of selection given by *s*(*t,N*(*t*)). Similar to Eqs.20 in Uecker and Hermisson (2011), the birth and death rates in the current model are given by

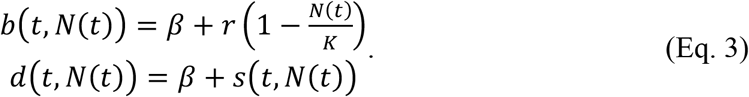

The parameter *β* is the constant birth and death rates of a population at carrying capacity if all individuals are at the optimum (no selection), and *r* is the intrinsic growth rate of the population (no selection). We define the average strength of selection as 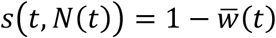, where 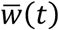 represents the mean fitness of the population at the *t*th generation: 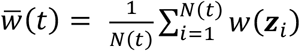. If the entire population is at the optimum, then 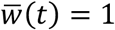 holds, and the population size approaches the carrying capacity, *K*. Deviations from the optimum, however, induce selection, which reduces the steady-state population size. In our model, 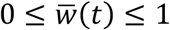 is always satisfied, and thus 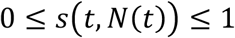 also holds, which ensures that the birth and death rates remain positive. Under the deterministic approximation, the total population size thus changes according to

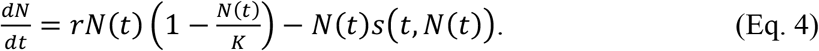

#### Extinction probability without adaptation

In the absence of adaptation due to de novo mutations, the population size is determined based on the initial mean fitness (i.e., mean phenotypic distance from the optimum), 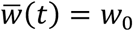, and approaches the following equilibrium:

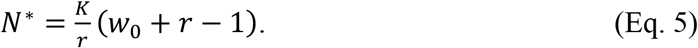

If *N*\* is close to 0, the population may become extinct by chance due to demographic stochasticity. To quantify the risk of extinction, we use the probability of loss in a general birth-death model starting from an initial state with *N*_0_ individuals;

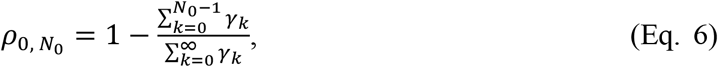

where 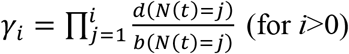 and *γ*_0_ = 1 (e.g., Otto and Day 2011).

#### Simulation methods

We performed simulations by initially sub-dividing the species into two finite populations, each comprising *K* individuals with no migration between them (i.e., allopatry). These populations are genetically identical initially, having just separated geographically.

To perform the evolutionary simulation with explicit population dynamics using SLiM3.3 (Haller and Messer 2019), the demography described in the birth-death process above needs to be approximated using the Wright-Fisher model with discrete generations. In a single population at time *t*, mating pairs are formed by randomly drawing individuals with replacement according to their relative fitness, and each pair produces a single offspring. This process is repeated *N*(*t* + 1) times, where *N*(*t* + 1) is the total number of offspring in the next generation, *t*+1. To determine *N*(*t* + 1), we consider the following stochastic differential equation to calculate the expected number of offspring in the next generation: Δ*N* = *M*(*N*(*t*))Δ*t* + *V*(*N*(*t*))Δ*ω*, where Δ*ω* is a random variable with mean 0 and variance equal to Δ*t*. The stochastic dynamics can be handled by considering the systematic change 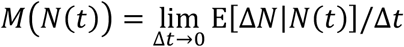 and the variance generation rate 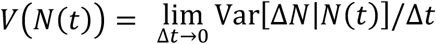 as follows:

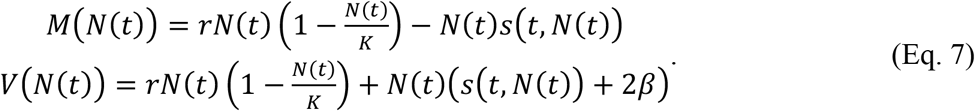

Thus, we can calculate the total number of offspring in the next generation: *N*(*t* + 1) = *N*(*t*) + Δ*N* with Δ*t* set to one.

We followed the populations for 4000 generations, unless otherwise stated. For the analysis of fitness reduction in hybrids, we focus on simulations in which the final fitness of both parental populations is higher than 0.95 to concentrate on those populations that had successfully adapted and avoid asymmetries in parental fitness that can affect hybrid fitness (i.e., these were considered to be still at risk of extinction and not included, see Table S1 for the number of simulations excluded).

#### Parallelism

Two parental populations on independent evolutionary paths may undergo adaptation by utilizing some common genetic loci. To quantify the degree of parallel evolution between two populations, we define “parallelism” as the fraction of the distance from the ancestral population to the optimum spanned by shared mutations (Figure 1c). At the end of each simulation, for alleles with a frequency of 95% or more in each population, we calculate parallelism. In the limit of the infinite-sites model, it is impossible to share the exact same mutation between populations, and thus parallelism becomes zero. However, if two populations complete their adaptation with exactly the same set of genetic variants (and the same phenotypic values), parallelism is 1. In rare cases, parallelism can be negative, which means that only maladaptive mutations (those that move away from the adaptive peak) are shared among the populations.

## Results

Under the parameters assumed (Table 1), the mean fitness of the population in the 1st generation is *w*_0_ ≈ 0.018, with an equilibrium population size (Eq.5) of *N*\* = 36 when *r* = 1. At such small population sizes, stochastic extinction is likely to occur, which was frequently observed when *r* was roughly lower than 1 (Figure S1). A rapid decrease in population size can, however, be followed by evolutionary rescue, if sufficiently large adaptive mutations arise soon enough (Figure S2). As expected, the population is more likely to be rescued when there are more sites under selection (Figure S1). Hereafter, we consider three scenarios with three different extinction risks: high extinction risk (*r*=0.8), moderate extinction risk (*r*=1.0), and low extinction risk (*r*=1.4).

As expected, populations at higher risk of extinction fixed larger-effect adaptive mutations, given that the population survived (Figure 2a). With a large number of mutable sites within the genome (*L*=10000, which we consider to be large and close to an “infinite-sites model”), we find that parallelism is almost zero except for a small fraction of parallelly evolved simulations (Table S1), and the fixation of larger effect mutations leads to greater reproductive isolation in populations that face a higher extinction risk, consistent with the phoenix hypothesis (Figure 2e).

**Figure 2.**
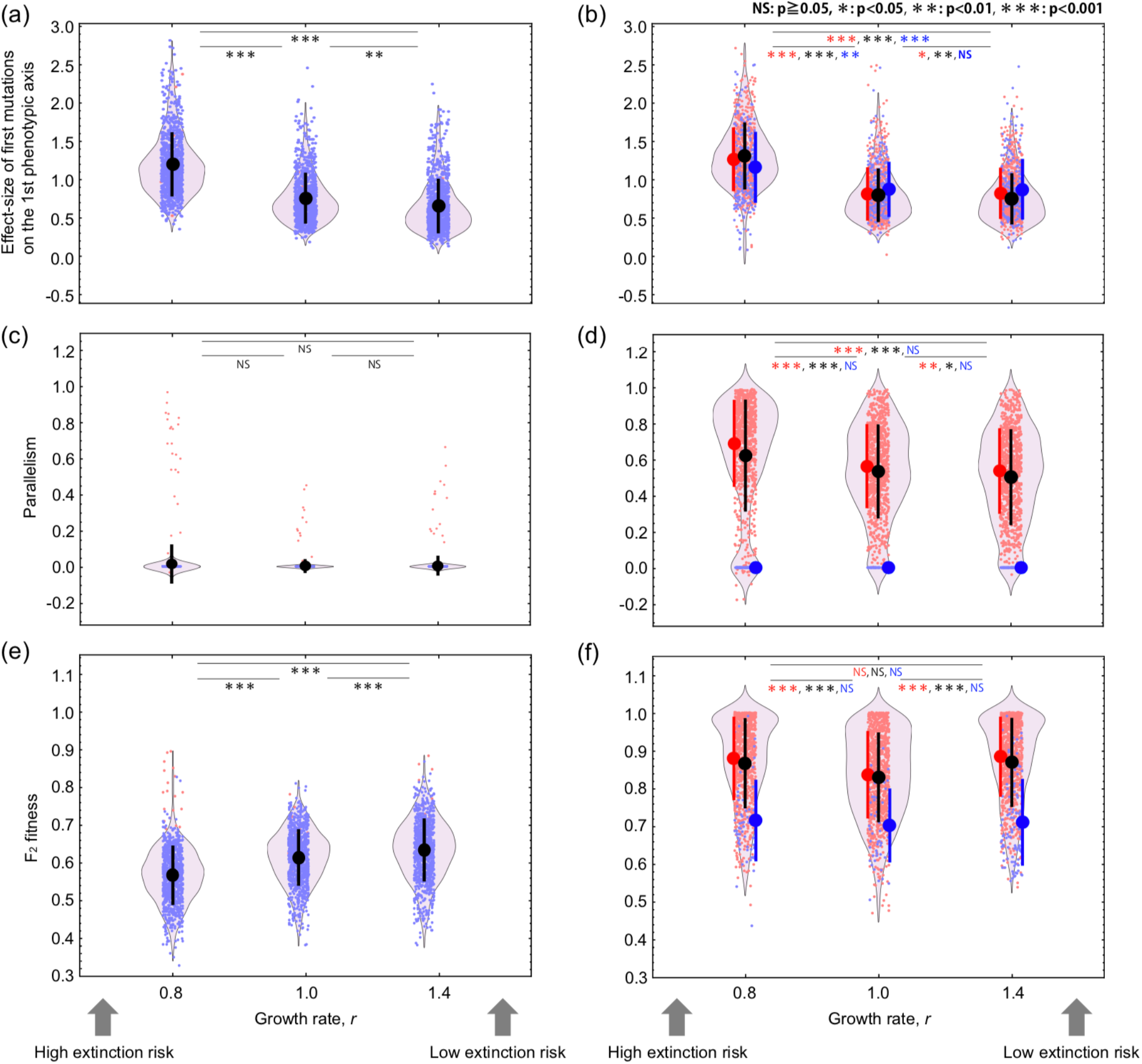
Characteristics of mutations that accumulate during the process of adaptation and the reduction of F_2_ fitness (left panels for *L*=10000 and right panels for *L*=50). Surviving populations at higher risk of extinction are more likely to adapt via large-effect mutations on the 1st phenotypic axis (the focal trait under environmental change) (a, b). With a large number of mutable sites within the genome, parallel adaptation rarely occurs (c). However, when the number of loci available for adaptation is finite, parallel mutations are more likely under a high extinction risk (d). F_2_ fitness decreases monotonically as extinction risk increases (e), which is expected in the Phoenix hypothesis. In the case of *L*=50, the net result is that populations facing an intermediate risk of extinction exhibit to lowest F_2_ hybrid fitness (f), consistent with a modified version of the phoenix hypothesis. The black dots and background violin plots represent the mean ± SD calculated from 1000 simulations. The 1000 simulations were divided into cases with zero and non-zero parallelism values, shown in blue and red, respectively. Likewise, solid points and lines indicate mean ± SD in right panels. The small dots are the values for each simulation run, and for the F_2_ fitness value, it is the average value of 2000 F_2_ hybrid individuals created. Kruskal-Wallis test is used. Parameters used are *L*=50 and *c*=0.

While such large-effect mutations are expected to increase the extent of reproductive isolation among populations, with a limited number of loci available for adaptation (*L*=50), the overall pattern of parallelism also becomes higher as extinction risk increases (Figure 2d), because surviving populations are more likely to re-use the same “rescue” mutations. Because of these contrasting effects, we find that hybrid fitness is lowest for populations at intermediate risk of extinction (Figure 2f), with different large-effect mutations incorporated in different populations. By contrast, when there is a low extinction risk, more mutations that accumulate during adaptation are unique, but those mutations are relatively small-effect and do not promote speciation. Therefore, moderate extinction risk results in the highest possibility of speciation due to the balance between mutation-size effect and mutation uniqueness. This pattern supports a modified version of the phoenix hypothesis that predicts a greater degree of reproductive isolation among populations experiencing a high extinction risk, but not so high that rescue frequently requires the same mutation(s).

We next asked if these results were robust to changing assumptions. First, we increased the number of sites (from *L=50* to 100). Similar results were obtained, except that parallelism did not display a monotonic decline with increasing probability of population survival (i.e., with increasing *r*). Specifically, populations at low and at high extinction risk showed significantly more parallelism than those populations at moderate risk of extinction (Figure S3, focusing on red points where some parallelism was observed). The parallelism observed at higher growth rates (*r*) was only slightly higher and likely reflects the maintenance of more genetic variation when the population size remained larger, favoring the same highly beneficial mutations especially when the number of loci (and hence the mutation rate) was greater. Consistent with this interpretation, fewer mutations fixed during the course of adaptation (Figure S4) when the extinction rate was high (fixing fewer and larger mutations) and when it was low (higher competition for the most beneficial mutations), compared to populations at moderate extinction risk.

We also explored changing the FGM to relax the assumption of complete pleiotropy, i.e., that each mutation affects each phenotype independently. Specifically, we allowed mutations to have a primary effect on a specific axis (no pleiotropy with probability 1-*c*), as well as a randomly drawn mutational effect with complete pleiotropy. With moderate pleiotropy (*c*=0.5), we again observed that a moderate extinction risk results in more extensive reproductive isolation (Figure S3 for *L*=100 and Figure S5 for *L*=50). The number of fixed mutations during adaptation also showed qualitatively similar results (Figure S4). These results support a modified version of the phoenix hypothesis, whether reproductive isolation results primarily from pleiotropic side-effects (*c*=0) or from overshooting effects (*c*=0.5). Furthermore, with no pleiotropy (*c*=1), the average degree of parallel evolution between two populations increases under any extinction risk (Figure S5d), and we observed the pattern of F_2_ fitness reduction as expected by the phoenix hypothesis (Figure S5f). In summary, whether a modified or the original version of the phoenix hypothesis is supported can vary depending on the genetic architecture underlying adaptation, including the number of loci and the degree of pleiotropy.

## Discussion

Allopatric speciation cannot be observed unless geographically isolated populations persist for the thousands or millions of years required for reproductive isolation to build (Futuyma 1987). During this period, environmental change can reduce the size of populations, increasing the likelihood of extinction due to demographic stochasticity (Lande 1993). In extreme cases, populations may be so maladapted that they decline in size to extinction unless beneficial mutations arise and lead to evolutionary rescue (Bell and Gonzalez 2009). In this study, we argue that populations that persist through such periods of rapid adaptation are more likely to accumulate the large-effect genetic changes likely to cause reproductive isolation. We call this process the phoenix hypothesis of speciation, because new species arise from the near extinction of populations that have undergone evolutionary rescue.

Reproductive isolation, as measured by a reduction in fitness of hybrids, is strongest when adaptation involves different large-effect mutations in different parental populations. Using FGM with the assumption of an infinite-sites model and constant population size, Yamaguchi and Otto (2020) showed that large-effect mutations can promote strong reproductive isolation even when populations are adapting to similar environmental changes. This previous work did not, however, consider population dynamics explicitly, and so did not explore the risk of extinction. Here we find an increase in the fixation of large-effect adaptive mutations in populations at high extinction risk and a greater reproductive isolation among those populations. Thus, the phoenix hypothesis was supported under the infinite-sites model. The study by Yamaguchi and Otto (2020) also did not consider the potential for parallel adaptation, which is high when few mutations can rescue a population (Figure 1). Parallel genetic evolution increases with the strength of natural selection and is particularly likely to occur for genes with large phenotypic effects (MacPherson and Nuismer 2017). We find that populations that persist despite a high risk of extinction are more likely to lead to accumulate large-effect adaptations, which more often generate reproductive isolation, but these adaptations are also more likely to be parallel, reducing the amount of reproductive isolation (Figure 2). Given these contrasting effects, we find that populations at an intermediate extinction risk develop the most reproductive isolation (Figure 2).

Because the magnitude of these contrasting effects depends on the parameters, the precise conditions most favorable to the phoenix hypothesis may not be as simple as illustrated in Figure 1ab. For example, we have observed non-monotonic trends for both parallelism (Figure S3) and the number of fixed mutations at the end of adaptation (Figure S4), results that might reflect stronger competitive replacement when many alternative mutations co-occur in large populations.

Changing population size is a key feature of a related model of speciation. Founder effect speciation considers the case where a small number of individuals colonize a new site, with genetic drift causing considerable genetic differentiation from the ancestral population (Mayr 1954; Templeton 1980). The founder effect can cause a rapid peak shift through an extreme bottleneck, but this differs substantially from our model, which focuses on adaptation to a single peak, with environmental change driving genetic change rather than drift.

Although we mainly focused on large-effect beneficial mutations and adaptation, fixation of weakly deleterious mutations by drift is likely to occur in a small population given enough time and a sufficiently high mutation rate. Deleterious mutations can also contribute to hybrid fitness breakdown because a certain fraction of the progeny will be expected to carry more deleterious mutations than either parent (Shpak 2005; Turelli et al. 2001). On the other hand, masking of these mutations in diploid hybrids can offset reproductive isolation, leading to hybrid vigor (Butlin 2005; MacPherson et al. 2022). Although we did not include unconditionally deleterious mutations, there is a chance that linked deleterious alleles hitchhike to fixation along with beneficial rescuing alleles (Hartfield and Otto 2011; Otto 2004). Such “undesirable hitchhikers” could also contribute to fitness differences between populations. Future work that includes such hitchhiking would be valuable to determine the extent to which the fixation of different deleterious mutations in different parental populations facilitates introgression and counteracts the phoenix hypothesis.

It would also be worthwhile exploring the effect of standing genetic variation or migration in future studies. Standing genetic variation enables the parental populations to incorporate the same alleles when the environment changes, increasing parallelism (Montejo-Kovacevich et al. 2021; Thompson et al. 2019). Similarly, migration between parental populations allows the spread and fixation of the same favorable alleles. A comparative genome study of nine- and three-spined stickleback populations revealed that small effective population sizes and restricted gene flow limit the potential for parallel local adaptation (Fang et al. 2021). Furthermore, the history of a population can impact reproductive isolation, with genetic surfing as the range of a population leading to “allele surfing” to high frequency of deleterious recessive alleles, which again facilitates introgression when a secondary contact occurs (MacPherson et al. 2022).

## Conclusion

The phoenix hypothesis predicts that populations are more likely to speciate if they have faced a history of environmental change so challenging that rapid adaptation was required to avoid extinction. While this hypothesis predicts a coupling between extinction risk and speciation, we emphasize that the new species formed by this process are not equivalent to the many lineages lost to extinction. Each lineage lost carries with it a long history of genetic adaptations. It has been estimated that new species arise, on average, at a rate of once per 2 MY (Hedges et al. 2015), and extinction eliminates all of the adaptations that allowed that species to survive and thrive since its last common ancestor with extant species. By contrast, newly formed species under the phoenix hypothesis can have minimal genetic differences, which makes them more prone to species collapse through introgression or competitive exclusion in the future (Seehausen et al. 2008).

With the accumulation of studies highlighting the role of large-effect mutations and adaptation, we find that rapid environmental change typically facilitates the evolution of reproductive isolation despite the higher extinction risk. Those lucky few populations that survive severe environmental change by evolving might, like the mythical phoenix, rise from the ashes reborn, taking the first baby steps to form a new species.

## Author contributions

RY and SPO designed the study. RY and BW developed the computer simulation code and RY conducted analyses in consultation with SPO. RY, BW and SPO wrote the manuscript.

## Data accessibility

All code necessary to repeat the analysis described in this study have been made available. SLiM source codes of our model for speciation dynamics will be hosted on Dryad Digital Repository upon acceptance. There are no data to be archived.

## Acknowledgements

This work was supported by funding from the JSPS KAKENHI (21K15160 and Overseas Research Fellowships to RY) and NSERC Discovery Grant RGPIN-2022-03726 to SPO.

## Online supplementary material

**Table S1.** Number of simulations for each parameter set we explored in this study. Of the 1000 successful simulations, we then show the average parental fitness and average F_2_ hybrid fitness.

| Parameters |  |  | Number of simulations |  |  |  | Fitnesses |  |
| --- | --- | --- | --- | --- | --- | --- | --- | --- |
| Genome size,<br>*L* | Degree of pleiotropic<br>side effect, *c* | Intrinsic growth<br>rate, *r* | Total | Excluded | Succeeded | Zero / Non-zero<br>parallelism | Mean parental fitness | Mean hybrid<br>fitness |
| 50 | 0.0 | 0.8 | 849658 | 79 | 1000 | 99/901 | 0.982223 | 0.868387 |
|  |  | 1.0 | 7775 | 20 | 1000 | 52/948 | 0.981885 | 0.830811 |
|  |  | 1.4 | 1054 | 54 | 1000 | 64/936 | 0.981568 | 0.870354 |
|  | 0.5 | 0.8 | 794199 | 83 | 1000 | 117/883 | 0.982362 | 0.876951 |
|  |  | 1.0 | 7672 | 91 | 1000 | 65/935 | 0.981734 | 0.827598 |
|  |  | 1.4 | 1045 | 45 | 1000 | 56/944 | 0.98143 | 0.859796 |
|  | 1.0 | 0.8 | 123737 | 67 | 1000 | 214/786 | 0.987939 | 0.918073 |
|  |  | 1.0 | 9603 | 37 | 1000 | 145/855 | 0.987396 | 0.933892 |
|  |  | 1.4 | 1046 | 46 | 1000 | 125/875 | 0.987207 | 0.951687 |
| 100 | 0.0 | 0.8 | 282259 | 70 | 1000 | 278/722 | 0.976815 | 0.778164 |
|  |  | 1.0 | 3824 | 61 | 1000 | 238/762 | 0.976829 | 0.729642 |
|  |  | 1.4 | 1099 | 99 | 1000 | 233/767 | 0.976933 | 0.759357 |
|  | 0.5 | 0.8 | 239263 | 24 | 1000 | 262/738 | 0.979541 | 0.782905 |
|  |  | 1.0 | 4010 | 42 | 1000 | 224/776 | 0.976762 | 0.730369 |
|  |  | 1.4 | 1084 | 84 | 1000 | 221/779 | 0.977031 | 0.761832 |
| 10000 | 0.0 | 0.8 | 228634 | 51 | 1000 | 975/25 | 0.981088 | 0.562212 |
|  |  | 1.0 | 3385 | 39 | 1000 | 986/14 | 0.984419 | 0.609702 |
|  |  | 1.4 | 1027 | 27 | 1000 | 984/16 | 0.98477 | 0.629868 |

**Figure S1.**
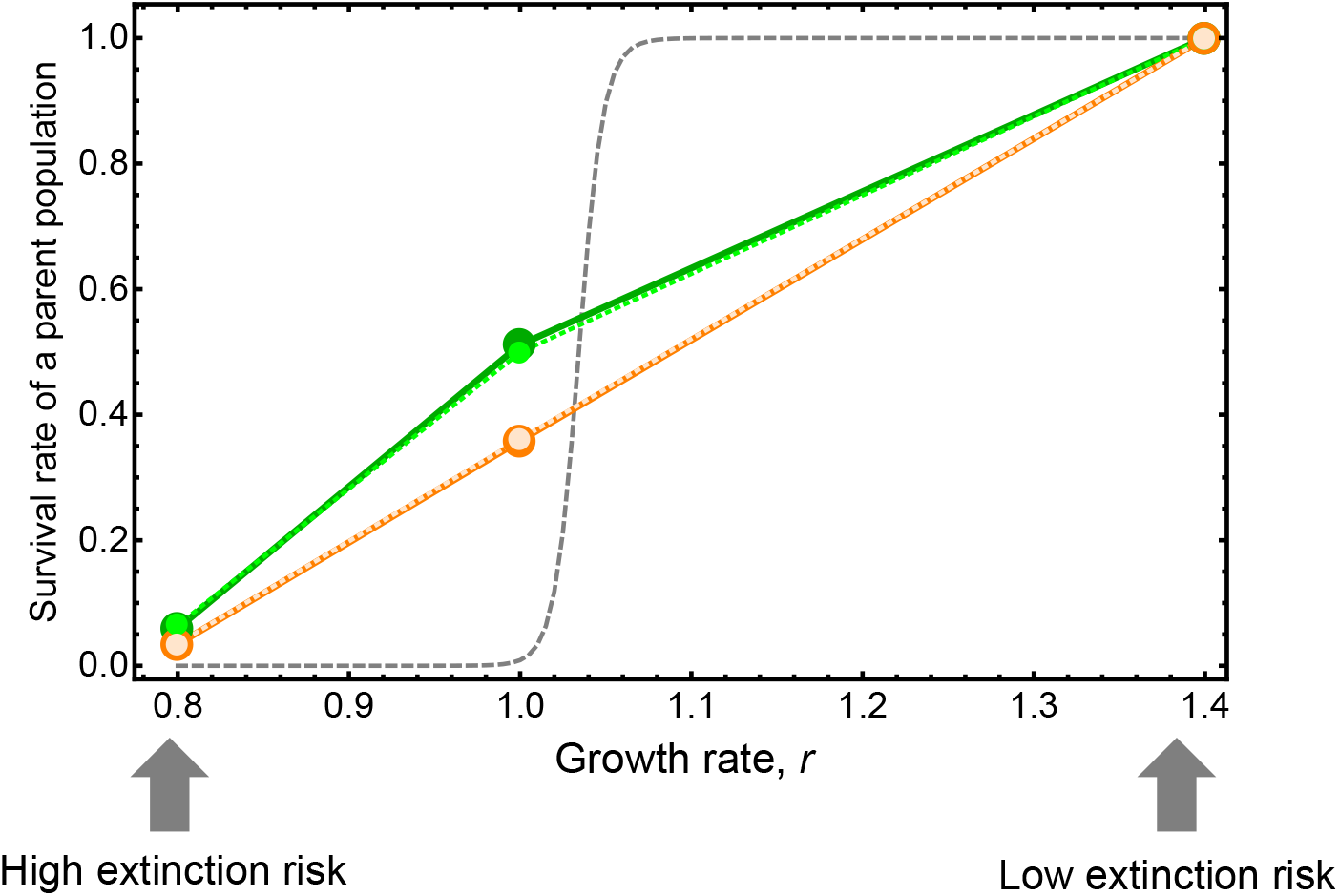
The survival rate of a single parent population against different intrinsic growth rates. Grey dashed line is calculated based on Eq. (6) assuming an initial population size of 200, and thus the survival rate corresponds to the situation without de novo mutations. As the growth rate increases, survival rate of the population rises from near zero (extinction without adaptation) to near one (low extinction risk). With mutations, survival probabilities rise due to evolutionary rescue. Solid lines and larger dots are results of *c*=0 (orange: *L*=50, green: *L*=100). Dotted lines and smaller dots are results of *c*=0.5 (lighter orange: *L*=50, lighter green: *L*=100). All other parameters are as in Figure 2.

**Figure S2.**
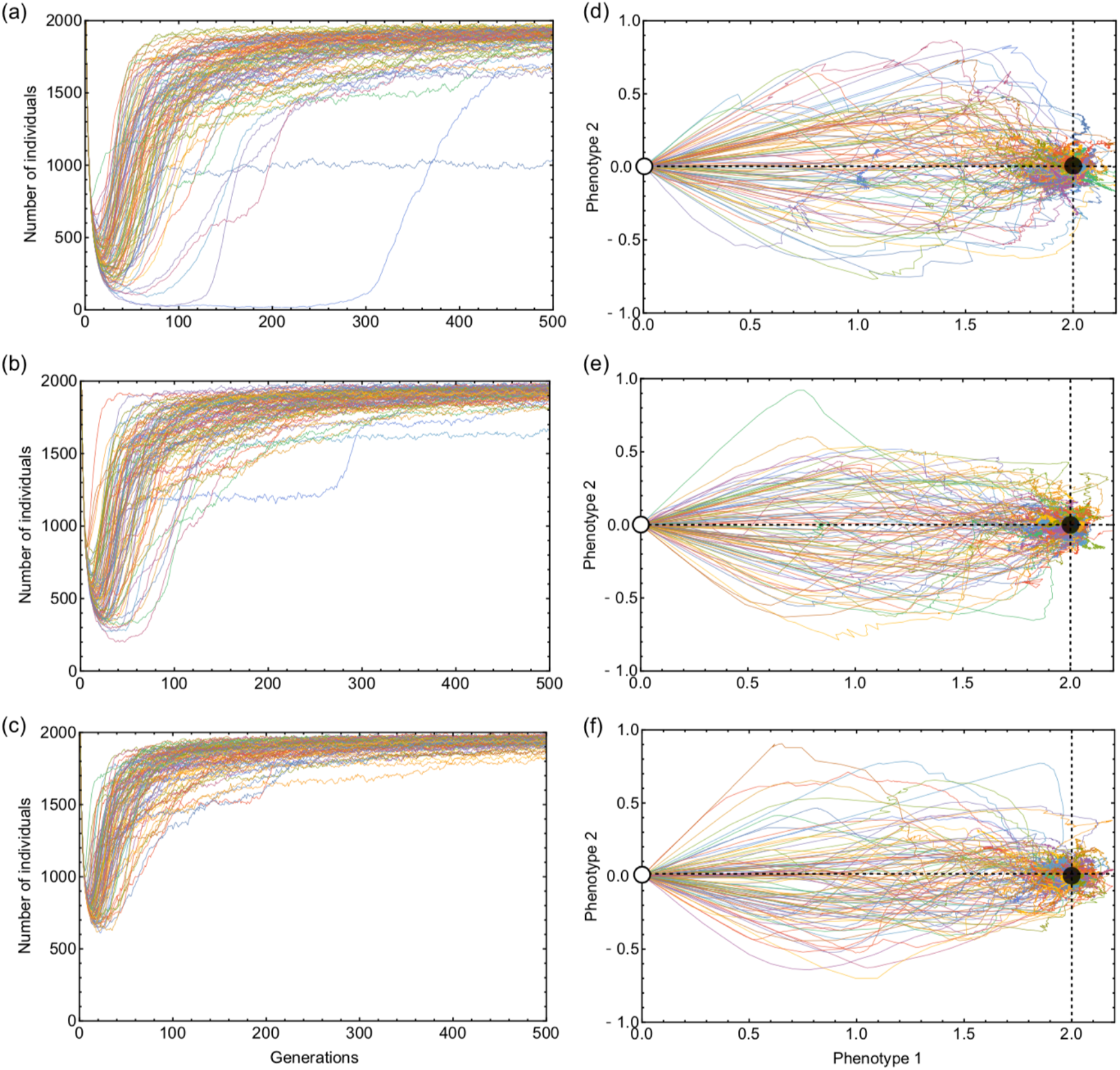
Example trajectories of population size (a,b,c) and phenotypes in two-dimensional space (d,e,f) during adaptation under different extinction risk: (a,d) high extinction risk *r*=0.8, (b,e) moderate extinction risk *r*=1.0, and (c,f) low extinction risk *r*=1.4. In right-hand side panels, white and black dots represent the initial and optimum phenotype, respectively. In all panels, 100 simulations were run for the parameter set as in the case of *L*=50 in Figure 2.

**Figure S3.**
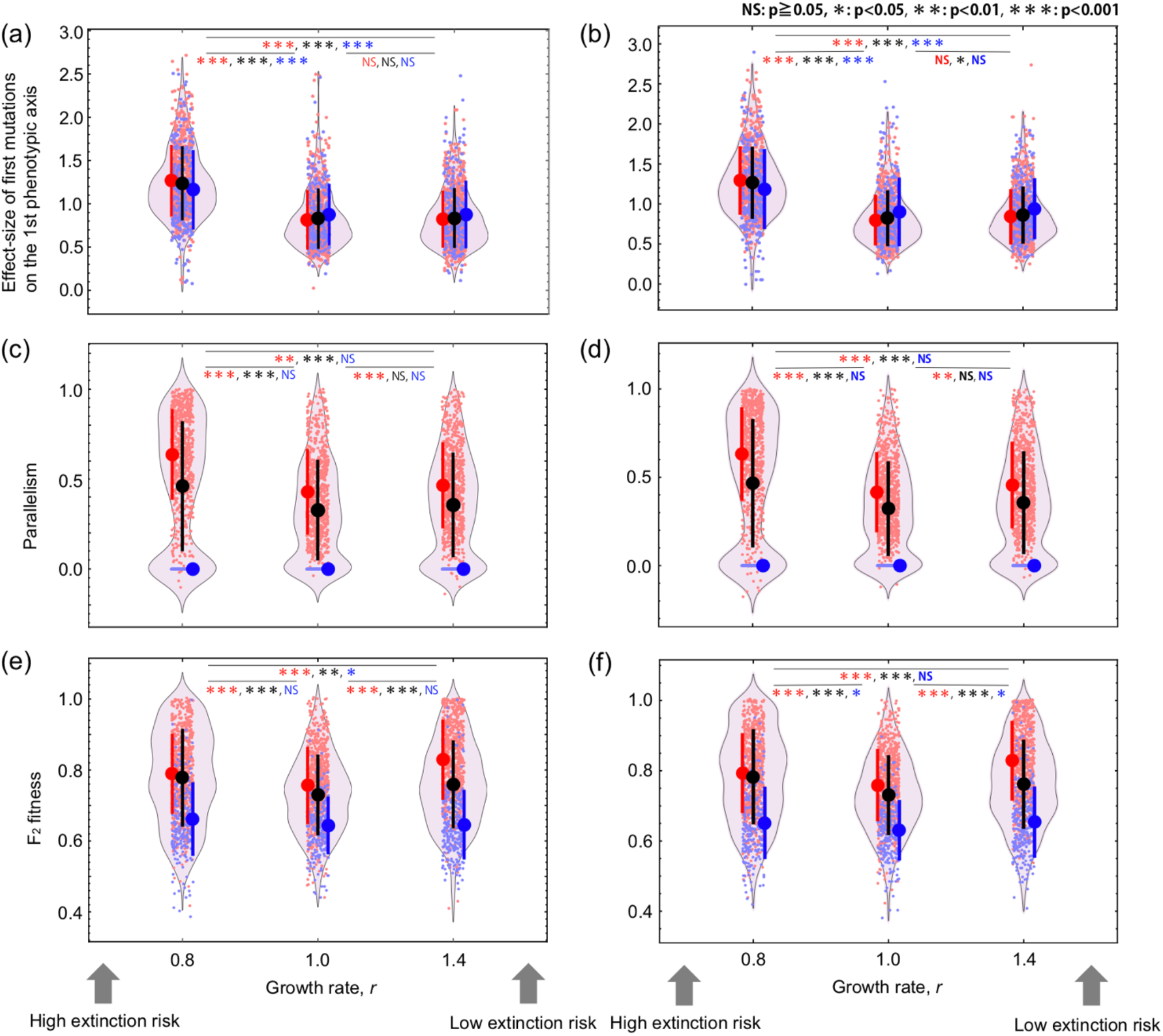
Characteristics of mutations that accumulate during the process of adaptation and the reduction of F_2_ fitness in the case of *L*=100 (left panels for *c*=0.0 and right panels for *c*=0.5). Surviving populations at higher risk of extinction are more likely to adapt via large-effect mutations on the 1st phenotypic axis (the focal trait under environmental change) (a, b) but to do so in a parallel manner as other populations (c, d). The net result is that populations facing an intermediate risk of extinction exhibit to lowest F_2_ hybrid fitness (e, f), consistent with the phoenix hypothesis. The black dots and background violin plots represent the mean ± SD calculated from 1000 simulations. The 1000 simulations were divided into cases with zero and non-zero parallelism values, shown in blue and red, respectively. Likewise, solid points and lines indicate mean ± SD. The small dots are the values for each simulation run, and for the F_2_ fitness value, it is the average value of 2000 F_2_ hybrid individuals created. Kruskal-Wallis test is used. Parameters used are the same as Figure 2.

**Figure S4.**
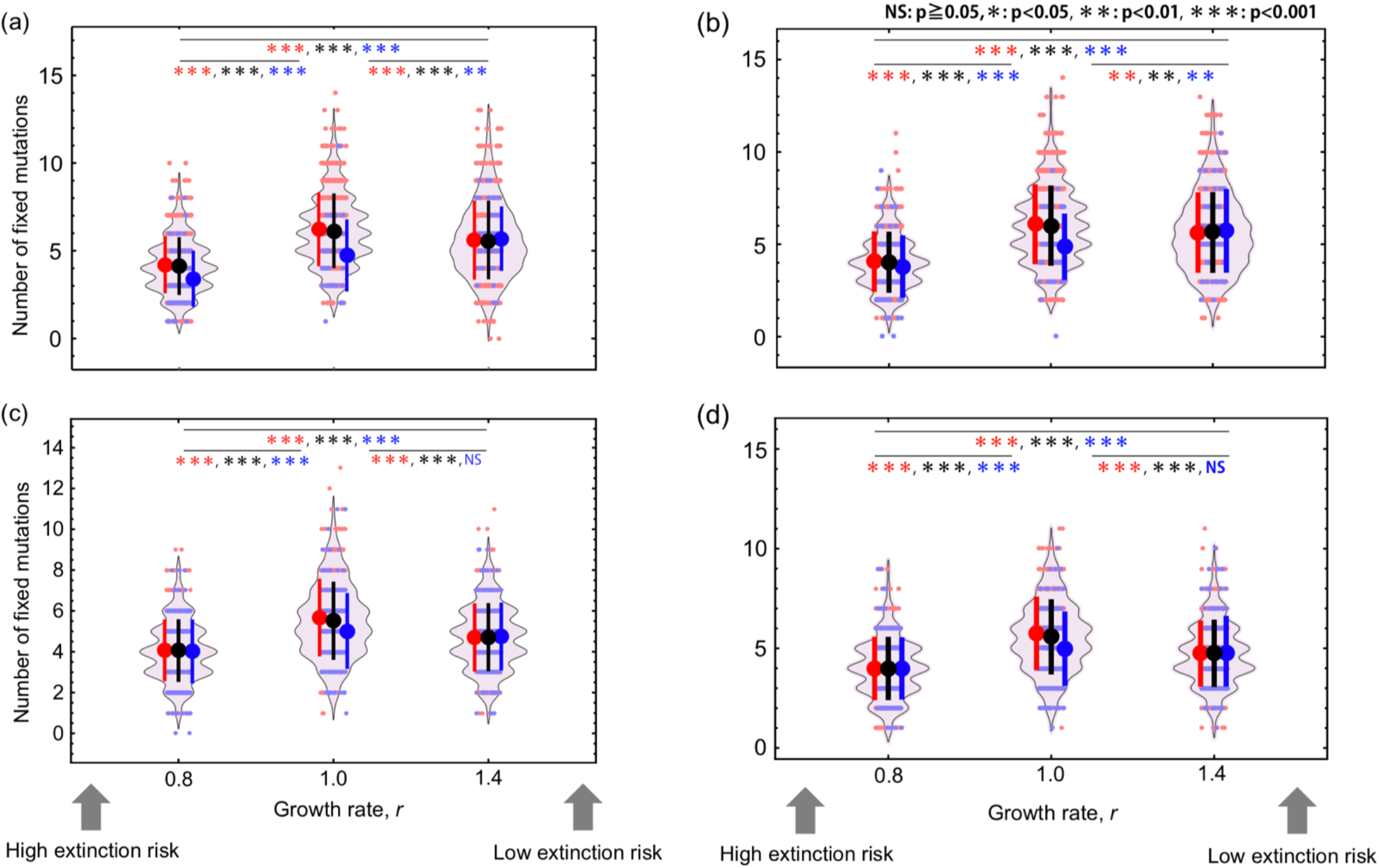
The number of fixed mutations at the end of each simulation. The upper panels (a, b) show the results of *L*=50 (left panel: *c*=0.0, and right panel: *c*=0.5), and the lower panels (c, d) show the results of *L*=100 (left panel: *c*=0.0, and right panel: *c*=0.5). An intermediate extinction risk maximizes the number except for the case of zero-parallelism of L=100. The black dots and background violin plots represent the mean ± SD calculated from 1000 simulations. The 1000 simulations were divided into cases with zero and non-zero parallelism values, shown in blue and red, respectively. Likewise, solid points and lines indicate mean ± SD. The small dots are the values for each simulation run. Kruskal-Wallis test is used. All parameters are as in Figure 2.

**Figure S5.**
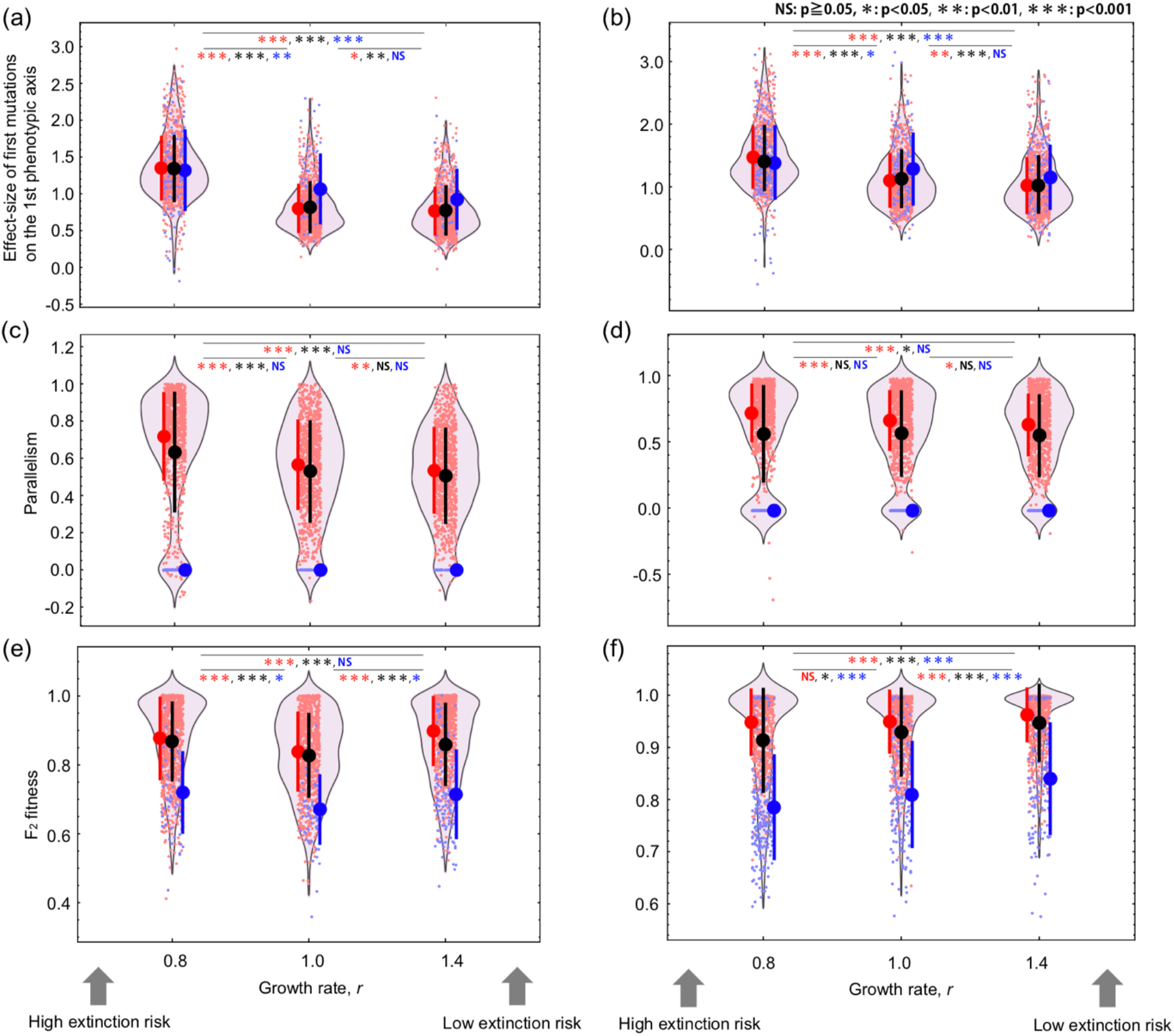
Characteristics of mutations that accumulate during the process of adaptation and the reduction of F_2_ fitness in the case of *L*=50 (left panels for *c*=0.5 and right panels for *c*=1.0). The black dots and background violin plots represent the mean ± SD calculated from 1000 simulations. The 1000 simulations were divided into cases with zero and non-zero parallelism values, shown in blue and red, respectively. Likewise, solid points and lines indicate mean ± SD. The small dots are the values for each simulation run, and for the F_2_ fitness value, it is the average value of 2000 F_2_ hybrid individuals created. Kruskal-Wallis test is used. Parameters used are the same as Figure 2.

